# Capillary-based Subcellular Sampling Uncovers the Stress Granule Proteome in Single Cells

**DOI:** 10.64898/2026.05.11.724230

**Authors:** Claire Davison, Nicolas Locker, Mariana Marques, Sam Kelly, Emily Relton, Tanavi Sharma, Emily Fraser, Pedro Aragon Fernandez, Erwin Schoof, Mia Petersen, Jordan Pascoe, Kathryn S. Lilley, Sneha Pinto, Matt Spick, Melanie Bailey

## Abstract

Many diseases arise from dysfunction within specific organelles or biomolecular condensates, highlighting the value of analysing proteins at subcellular resolution to uncover new biological mechanisms. We report a novel capillary-based subcellular sampling workflow coupled with liquid chromatography-mass spectrometry (LC-MS) for proteomic analysis of defined subcellular regions of individual cells. We applied this methodology to stress granules (SGs), membrane-less biomolecular condensates that form in response to cellular stress (including viral infection), and are implicated in infection, neuropathology and cancer. Comprehensive characterisation of SG protein composition remains limited by technical challenges associated with bulk purification, including loss of spatial context, dynamic behaviour and contamination from cytosolic material. Using our novel method, we identified a high-confidence set of 405 SG-associated proteins, including 46 established SG residents alongside numerous previously unreported candidates. Functional enrichment analysis revealed pathways consistent with known SG biology, while comparison with an independent cytosolic proteome dataset demonstrated minimal overlap, supporting the specificity of the sampling strategy. Selected novel SG protein candidates (AHNAK2, DDX39B, NUDT1 and FKBP2) were validated using immunofluorescence microscopy. These findings establish capillary-based subcellular sampling as a viable approach for proteomic analysis of SGs with preserved spatial context and provide a framework for analysing other subcellular compartments.

**Table of contents:** We report an LC–MS-based capillary sampling workflow for proteomic analysis of subcellular structures within single cells. This methodology identified 405 high-confidence stress granule-associated proteins, including 46 previously established and numerous novel candidates. The approach demonstrated high specificity and preserved spatial context, expanding the capabilities of subcellular proteomics.

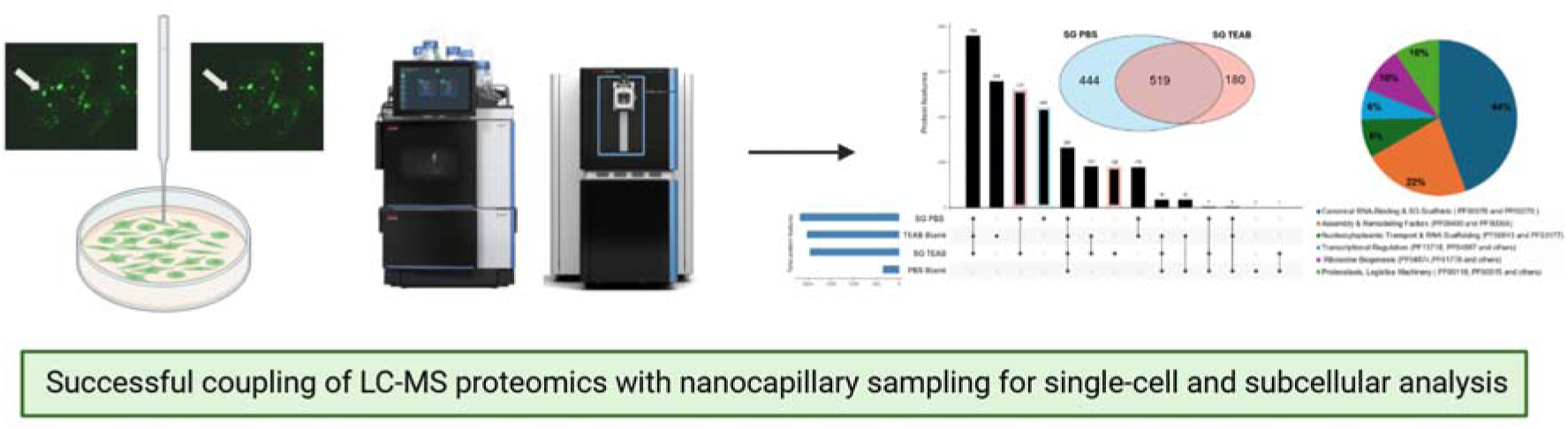

Figure made in Biorender.com.

## Introduction

Environmental and physiological stresses (e.g. viral infection, nutrient deprivation, hypoxia, heat, metabolic dysfunction) can profoundly affect a cell’s ability to maintain homeostasis. To ensure survival under these conditions, cells activate coordinated stress-response pathways that can reprogram transcription and translation within RNA-protein networks [1] [2]. In response to stress, membrane-less biomolecular condensates such as SGs concentrate RNAs and proteins via phase separation, creating dynamic signalling hubs that separate components from the cell milieu [3]. This allows cells to rapidly adjust processes and tune biochemical pathways following physiological and pathological triggers [4] [5]. Following the activation of the integrated stress response and inhibition of protein synthesis, stalled mRNAs accumulate in cytoplasmic RNP complexes, subsequently recognised by aggregation prone RNA-binding proteins (RBPs) and additional intrinsically disordered proteins [3]. The scaffold proteins G3BP1/2 then drive the condensation of SGs, bridging multivalent interactions between RNAs and proteins [6] [7]. Upon stress removal, the resolution of SGs releases mRNAs for translation to restore homeostasis.

Dysregulated SGs, often driven by mutation, may result in aberrant or persistence SG assembly and failure to clear aggregated proteins. Such dysregulation may be associated with diseases, including infection, neuropathology and cancer [8]. Furthermore, in response to different stresses, SGs have been proposed to adopt specific composition to tailor cellular outcomes, including death or survival [9], and selectively triage and condense resident proteins to drive specific cellular functions [9] [10]. For example, SGs induced in response to UV damage, compared to canonical arsenite-induced SGs, preferentially enrich for intronic mRNAs and the RNA helicase DHX9 but lack PKR. Their composition has been proposed to enable SGs to act as platforms for pathogen-associated molecular patterns (PAMPs), facilitating the removal of aberrant proteins or RNAs during UV damage [11]. Similarly, although this remains debated, in response to viral infection or double-stranded RNA stimulation, SGs have been proposed to be involved in innate immunity through the accumulation of innate immunity effectors including PKR, MDA5, RIG-I and OAS3 [12] [13]. Accumulation of these pattern recognition receptors (PRRs) in SGs may limit their interaction with dsRNA and thereby dampen the innate immune response [14]. Therefore determining which proteins accumulate within SGs under different conditions is crucial for understanding why these structures become dysfunctional in disease and for identifying potential targets for therapeutic intervention [15].

Defining SG protein composition using compositional approaches has deepened our understanding of SG biology and allowed us to characterise SG assembled in response to oxidative stress, endoplasmic reticulum (ER) stress or novel virus-induced SG-like foci [9] [16]. Despite these advances, obtaining an accurate profile of SG components remains technically challenging [17] [18]. SG-associated proteins typically constitute only a small fraction of the total cellular protein pool, making confident detection above the background signal difficult. Consequently, approaches based on proximity ligation may not discriminate between interactions occurring within granules or those mediated by the cytoplasmic fraction of the bait protein. SGs also possess a highly heterogeneous molecular composition (e.g. untranslated mRNAs, translation initiation factors, RNA-binding proteins, metabolic enzymes) which varies across different stressors, cell types and experimental conditions [17] [19] [20]. Moreover, intact isolation of individual SGs can be hindered due to their liquid-like biophysical properties, dynamic behaviour and lack of a membrane boundary [18]. Therefore, approaches based on differential centrifugation and affinity purification can result in the loss of individual components associated with the SG-dynamic shell structure.

Due to these technical challenges, bulk mass spectrometry methodologies are typically favoured over single-cell approaches for studying SG protein composition, despite the inherent loss of spatial resolution and information on cell-to-cell heterogeneity. These bulk approaches also have the disadvantage of extensive sample preparation steps [21] [22] [23] that can promote co-purification of non-specific cytosolic proteins [24] [25]. These limitations highlight the need for alternative analytical strategies that enable targeted, spatially resolved isolation of SGs prior to downstream analysis by mass spectrometry.

Capillary sampling has emerged as a powerful tool for isolating user-selected single cells under microscope observation prior to downstream mass spectrometry analysis, enabling targeted extraction of live cellular material whilst retaining spatial information. This sampling approach has been demonstrated for liquid chromatography (LC-MS) measurements in single whole cells, with successful applications in lipidomics [26] [27] [28] [29] [30] [31], metabolomics [32], and drug measurements [33]. Previous studies have also demonstrated the extraction of specific subcellular compartments, including mitochondria [34] [35] [36], lipid droplets [37] [38] and discrete nuclear and cytoplasmic fractions [39]. Together, these studies highlight the potential of capillary-based approaches for probing biological processes at subcellular resolution.

However, whilst effective for small-molecule analysis, capillary-based workflows remain considerably less established for proteomic profiling of small subcellular structures such as SGs. To date, existing single cell methodologies have predominantly employed microfluidic approaches [40] [41] [42] and laser capture microdissection [43] [44] for single cell capture prior to proteomic analysis. It is also possible to infer the protein content of sub-cellular organelles in bulk using sub-cellular fractionation [45]; however, this approach does not allow proteins to be probed from within a specific single cell. Single-cell and subcellular proteomics have also been achieved using capillary electrophoresis, however this has been demonstrated primarily in large embryonic cells, where sufficient material can be obtained for protein identification [46] [47]. To our knowledge, there are no examples of capillary sampling being coupled with LC-MS for proteomic analysis. SGs therefore represent a particularly challenging and unexplored test case for this.

To meet this challenge, we integrated capillary sampling with single-cell proteomics to enable spatially resolved single and subcellular analysis. In this workflow, live cell imaging is first used to identify structures of interest using fluorescent reporters, allowing selection of cells with desired phenotypes prior to sampling. Once the target compartment is located and confirmed, a fine glass capillary is positioned to isolate the specific region from the cell for downstream analysis. Here we apply this methodology to U2OS whole cells and SGs, enabling proteomic analysis of defined subcellular regions from individual cells. Using this approach, we identify established SG components for method validation while also revealing previously unreported candidates. Selected SG protein candidates are further supported by immunofluorescence localisation, confirming their recruitment to SGs and demonstrating the utility of this workflow for expanding our understanding of SG composition.

## Methods

### Cell culture and stress granule induction

U2OS G3BP1-GFP cell lines were grown in Dulbecco’s modified Eagle’s medium (DMEM) supplemented with 10% fetal bovine serum, 100 U/ml penicillin, 100 mg/ml streptomycin and 1% v/v L-glutamine, in a humidified incubator at 37°C and 5% CO_2_ environment.

For SG assembly, sodium arsenite (Sigma Aldrich) was diluted in H_2_O and used at a concentration of 0.5 mM for 30 min. Silvestrol (MedChemExpress) was resuspended in dimethyl sulfoxide (DMSO) and used at a concentration of 2 μM for 1 h.

### Single cell and stress granule sampling

Single cells and SGs were sampled using the Yokogawa SS2000 Single Cellome System (see **Figure 1**). Prior to sampling, untreated control cells and stressed cells were transferred into the built-in incubator of the sampling chamber to be maintained under conditions of 37 °C with 5% CO2 throughout the live-cell sampling procedure. Whole cell sampling was performed using 10 µm capillary tips under confocal imaging, while individual SGs were isolated using 3 µm capillary tips under fluorescence microscopy. To facilitate identification and targeted sampling of SGs, cells constitutively expressed the essential SG scaffolding protein G3BP1-green fluorescent protein (G3B1-GFP) as previously used [48] (excitation channel 405 nm nm and emission channel BP525/50 nm).

**Figure 1.**
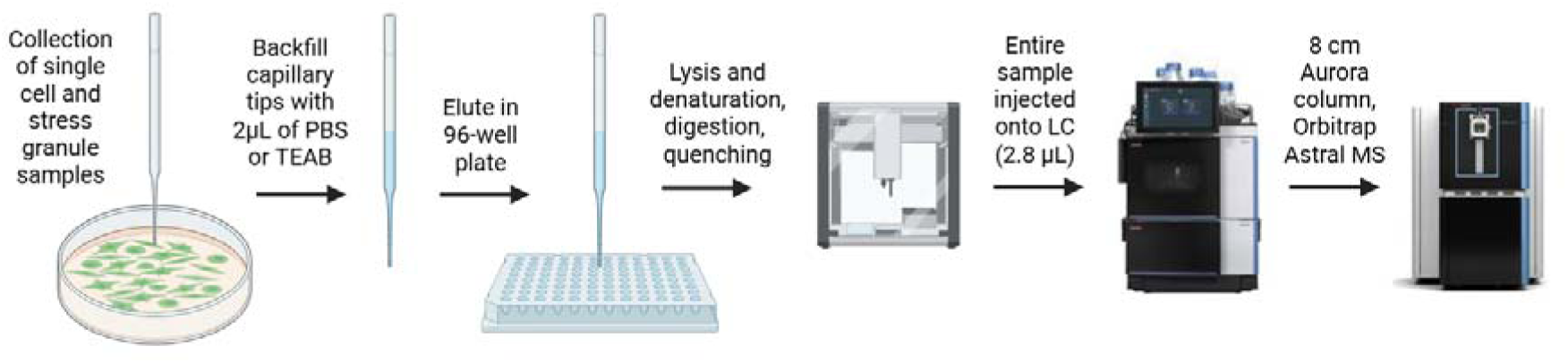
Proteomic workflow for capillary-based sampling and LC–MS analysis of single whole cells and SGs. Figure made in Biorender.com. Images of Vanquish Neo Ultra-High Performance Liquid Chromatography (UHPLC) and Orbitrap Astral mass spectrometer taken from *ThermoFisher.com*.

Sampling parameters were optimised separately for whole-cell and SG isolation to account for differences in size, puncture requirements and mechanical resistance. Whole-cell sampling employed higher sampling aspiration pressures of (-15.0) - (-14.7) kPa and post sampling pressures of (-0.8) - (-0.7) kPa, together with holding times of 500-1000 ms. In contrast, SG sampling was performed using reduced sampling aspiration pressures of (-25.9) – (-15) kPa and post sampling pressures of (-1.1) – (-0.9) kPa, but higher hold times of 5000 ms. For both whole cells and SGs, pre-pressures of 5.1-5.4 kPa were used. Blank samples were collected by sampling from a dish containing only either PBS or triethylammonium bicarbonate (TEAB; 80 mM, 0.2% n-dodecyl-β-D-maltoside) and stored as described for single cells.

### Nanocapillary elution and sample preparation

Following collection of single-cell and subcellular samples, capillary tips were backfilled with 2 µL of either PBS or TEAB. Samples were subsequently eluted from the capillary tips by manual pressure injector using a single-syringe infusion pump (KD Scientific) into low-bind 384-well plates. Eluted samples and corresponding blanks (comprising the elution solvent only, eluted through a capillary and into the 384 well plate) were immediately stored at -80°C and shipped on dry ice to the Technical University of Denmark for downstream proteomic analysis.

### Proteomic analysis by LC-MS

96-well plates containing the samples were briefly spun down, snap-frozen on dry ice for 5 minutes, and then heated at 95°C in a PCR machine (Applied Biosystems Veriti, 384-well) for an additional 5 minutes. Samples were then either subjected to tryptic digestion or stored at −80°C until further processing. Tryptic digestion of samples was carried out by adding 1 μL of Trypsin platinum mix (2 ng/μL in 100 mM TEAB, Promega) and incubation at 37°C overnight, followed by the addition of 1 μL of trifluoroacetic acid (TFA, 1 % (v/v)). All reagent dispensing was done using an I-DOT One instrument (Dispendix).

Peptides were separated using a Vanquish Neo Ultra-High Performance Liquid Chromatography (UHPLC) system operated in nano-flow mode. Samples were loaded at 60 µL/min under pressure-controlled conditions onto a PepMap C18 trap column (Thermo Fisher Scientific) and separated in a trap-and-elute configuration on an Aurora Series C18 analytical column (8 cm × 75 µm ID; IonOpticks). The analytical column temperature was maintained at 50 °C through all acquisitions. Chromatographic separation was performed using solvent A (water with 0.1% formic acid) and solvent B (80 % acetonitrile with 0.1% formic acid in water). Peptides were eluted at 100 nL/min using a short linear gradient from 4 to 35% B, followed by a high-organic wash at 99% B. The system was subsequently re-equilibrated to initial conditions prior to the next injection. The total LC method time was 10.1 min corresponding to 120 samples per day.

Peptides were analysed on an Orbitrap Astral mass spectrometer (Thermo Fisher Scientific) operated in positive ion mode with a nano-electrospray ionization (NSI) source. The spray voltage was set to 1.9 kV, and the ion transfer tube temperature was 240 °C. Field Asymmetric Ion Mobility Spectrometry (FAIMS) was enabled in standard resolution mode using a compensation voltage of −48 V. Data was acquired using a data-independent acquisition (DIA). Full MS scans were acquired in the Orbitrap at a resolution of 240,000 over an *m/z* range of 400–800, with an AGC target set to 500 % and a maximum injection time of 100 ms. DIA MS/MS scans were acquired using the Astral analyser over an *m/z* range of 150–2000 with a cycle time of 0.6 s, employing isolation windows spanning *m/z* 400–800. Fragmentation was performed by higher-energy collisional dissociation (HCD) using a normalised collision energy of 25%. All spectra were acquired in profile mode.

### Data processing and statistical analysis

For the mass spectrometry raw data analysis, DIA data were processed using DIA-NN v1.9.2 [49]. Raw files were searched in library-free mode with *in silico* spectral library generation enabled. Peptide identification was performed against a combined FASTA database comprising the reviewed *Homo sapiens* proteome (UniProt, February 2025). Trypsin specificity was assumed with cleavage at K/R residues, allowing up to one missed cleavage. The precursor *m/z* range was set to 400–800 *m/z* with charge states +1 to +4, and fragment ions were considered in the 200–1800 *m/z* range. Peptide lengths were restricted to 7–30 amino acids. N-terminal methionine excision was enabled. Oxidation of methionine (UniMod:35) was included as the only variable modification, with a maximum of one variable modification per peptide. False discovery rate (FDR) control was applied at 1 % at the precursor and protein levels. Quantification matrices for precursors and protein groups were exported for further analysis.

All further statistical analyses and data visualisation were performed in R (RStudio, version 2026.01.0). Unless otherwise specified, no global frequency-based filtering was applied to the proteomic dataset at the initial analysis stage. Frequency-based analyses were instead applied explicitly at later stages to assess reproducibility and consistency of protein detection across SG samples (as described in the **Results** section).

Functional association network and pathway enrichment analyses were performed using the STRING database (version 12.0, 2023) [50], which integrates known and predicted protein-protein interactions and functional annotations. Enrichment significance was assessed using false discovery rate (FDR)-corrected p-values based on the Benjamini-Hochberg procedure, as implemented in STRING. Pathway enrichment strength was additionally summarised using the STRING signal score, a composite metric that integrates enrichment magnitude (observed-to-expected ratio) with statistical significance to enable intuitive ranking of enriched functional terms.

Given that intrinsically disordered regions (IDRs) are a hallmark feature of SG proteins [51], a systematic structural analysis of our filtered dataset was performed. Sequence features were retrieved from UniProtKB, utilizing the MobiDB-lite consensus to ensure high-confidence mapping of disorder boundaries through multiple independent algorithms. To further characterize these disordered regions, we identified low-complexity regions (LCRs) and specific sequence enrichments using UniProt’s integrated compositional bias annotations (e.g. ‘low complexity’, ‘basic and acidic residues’), and also cross-referenced against the experimental “gold-standard” evidence from the DisProt database (Release 2026). By evaluating the dataset against manually curated and experimentally supported disordered regions independently of the computational LCR predictions, we were able to identify a high-confidence subset of proteins inclusive of diverse disordered architectures.

### Immunofluorescence validation

Cells were seeded on sterilised coverslips in a 24-well plate and treated as required (see **Cell culture and stress granule induction** section). Media was removed and fixed with 4% paraformaldehyde (PFA) in PBS for 20 min at RT. Fixation solution was removed, coverslips were rinsed with PBS and stored at 4 °C or processed immediately. Cells were permeabilized with 0.1% Triton X-100 (Sigma) in PBS for 5 min at room temperature (RT) and blocked with 0.5% bovine serum albumin (Fisher) in PBS for 1 h at RT. Cells were then incubated with primary antibody solution for 1 h at RT, washed 3 times with PBS and incubated with secondary antibody solution containing 0.2 μg/ml DAPI for 1 h at RT. Coverslips were then rinsed 3 times with PBS and mounted onto microscope slides with 5 μl Mowiol 4-88 (Sigma-Aldrich #81381). Cells were visualised and imaged with a Nikon Ti-Eclipse A1M Confocal Microscope using a 60X objective. Confocal images were analysed utilizing Image J. The primary antibodies used were rabbit anti-G3BP1 (1:600, Sigma), goat anti-eIF3η (1:600, Santa Cruz). Secondary antibodies used were goat anti-rabbit Alexa 488, donkey anti-goat Alexa 555, chicken anti-mouse 633, goat anti-mouse-IgG2 Alexa 568 and goat anti-mouse-IgG1 Alexa 488 (1:500, Invitrogen).

## Results

U2OS cells were used as benchmark system for prior compositional analysis of SG resident protein, and sodium arsenite was employed as positive control for SG assembly, as it induces eIF2α-dependent SGs through activation of the eIF2α kinase HRI [52]. A total of 5115 proteins were identified from the complete dataset containing the six sample types (PBS blank control, TEAB blank control, PBS eluted SGs, TEAB eluted SGs, stressed cells and untreated control cells). Of these, 5022 proteins were detected across SGs and whole cells under stressed and unstressed conditions. This broad proteomic coverage provided a foundation for assessing both whole-cell stress responses and SG–associated protein composition using the capillary-based single-cell workflow.

We first assessed whether capillary sampling could be coupled to the single-cell proteomic workflow to detect biologically meaningful changes associated with stress at the single cell level. To achieve this, whole stressed single-cells (n=10) were compared to untreated single-cell controls (n=13). Across this comparison, numerous proteins exhibited statistically significant differences in abundance between stressed and control conditions (see **Figure 2** and SI Table 1). Specifically, 75 proteins had increased abundance and 277 had decreased abundance, indicating a robust proteomic response to stress induction. These results demonstrate that the capillary-based sampling and LC-MS workflow is sensitive to cellular stress state and capable of resolving stress-associated proteomic differences.

**Figure 2.**
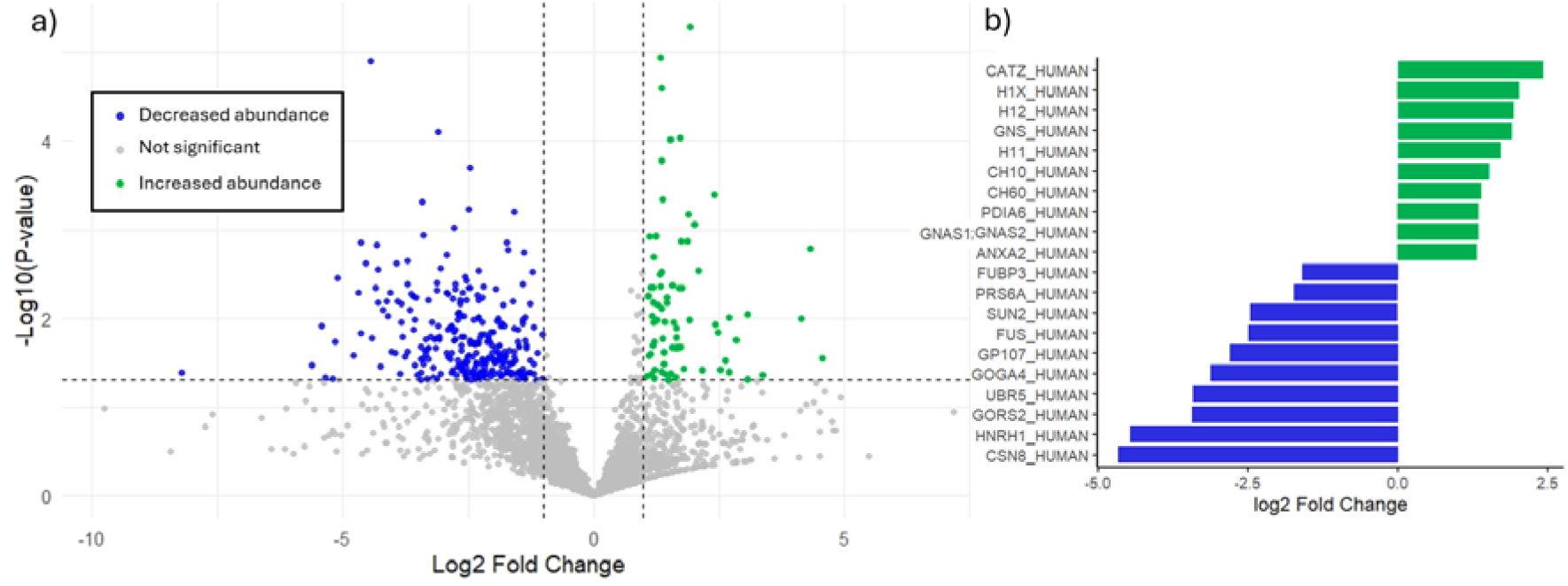
**a)** Volcano plot showing differential protein abundance between stressed and untreated whole cells sampled in PBS. Proteins were considered significantly differentially abundant at *p* < 0.05 and |log_₂_FC| > 1. Upregulated proteins (green) have log_₂_FC > 1 (higher in Whole Stressed Single-Cells), and downregulated proteins (blue) have log_₂_FC < −1 (higher in untreated single-cell controls); proteins not meeting these thresholds were considered not significant. **b)** Bar plot showing the top 10 increased abundance and top 10 decreased abundance proteins ranked by Benjamini–Hochberg false discovery rate (FDR)-adjusted p-value (padj). Bars indicate log2 fold change (log2FC) relative to the comparator condition; positive values denote increased abundance (green), and negative values denote decreased abundance (blue).

Having established that the workflow captured stress-dependent proteomic differences at the whole cell level, we next applied this approach to the targeted sampling of SGs. SGs were readily identified in live cells using a G3BP1 – green fluorescent protein reporter, enabling targeted sampling under fluorescent microscopy (see **Figure 3**). Time-resolved imaging confirmed selective puncture at the SG site, supporting the feasibility of capillary-based extraction of subcellular compartments for downstream analysis.

**Figure 3:**
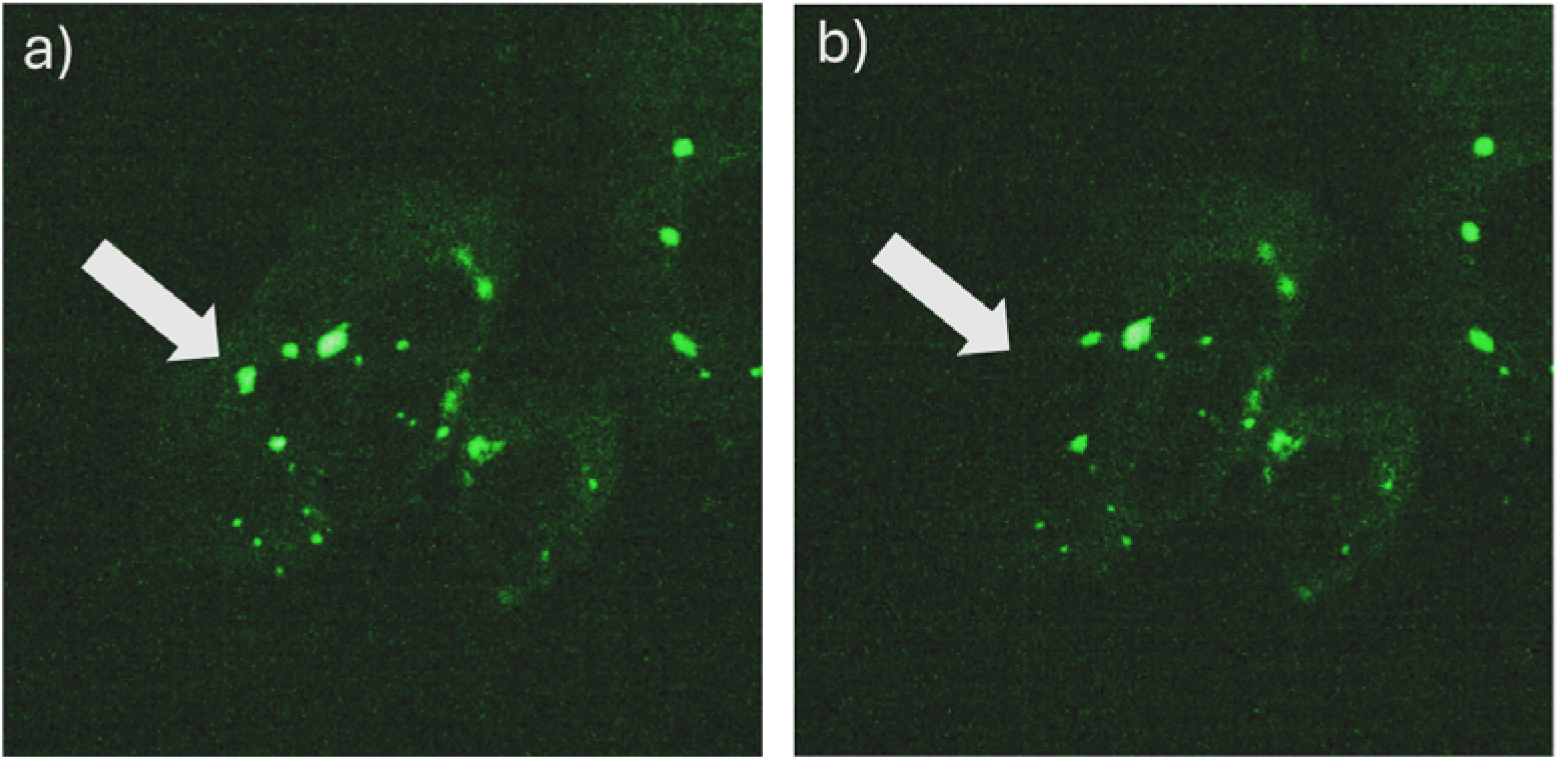
Fluorescent microscopy guided capillary sampling of SGs. SGs identified using G3BP1-GFP (green) **a)** prior to sampling, **b)** immediately following capillary puncture and uptake of SG into tip.

As part of the sampling strategy, SG samples were eluted using two solvent conditions prior to proteolysis and downstream processing, PBS (n=5) and TEAB (n=7), to generate parallel datasets of the SG proteome. Although TEAB is more suitable for downstream proteomic analysis by LC-MS, experimentally it led to bubbling and sample loss during elution from the capillary. In contract, PBS enabled reliable recovery of single cells without bubbling.

SG samples (each from a different cell) were subsequently analysed to define a high-confidence SG proteome from the full dataset of 5115 proteins. To minimise inclusion of proteins arising from contaminants on the capillaries, vials, solvents or mass spectrometry system, and to increase confidence in SG assignment, only proteins that satisfied three criteria were retained: detection in at least one PBS-eluted SG, detection in at least one TEAB-eluted SG, and absent from both PBS (n=6) and TEAB (n=5) blank media controls (see **Figure 4**). By requiring consistency across two independent elution conditions and exclusion from matched blanks, this approach reduced false-positive assignments and resulted in a final set of 519 features that were considered high-confidence SG-associated proteins (see **SI Table 2**).

**Figure 4.**
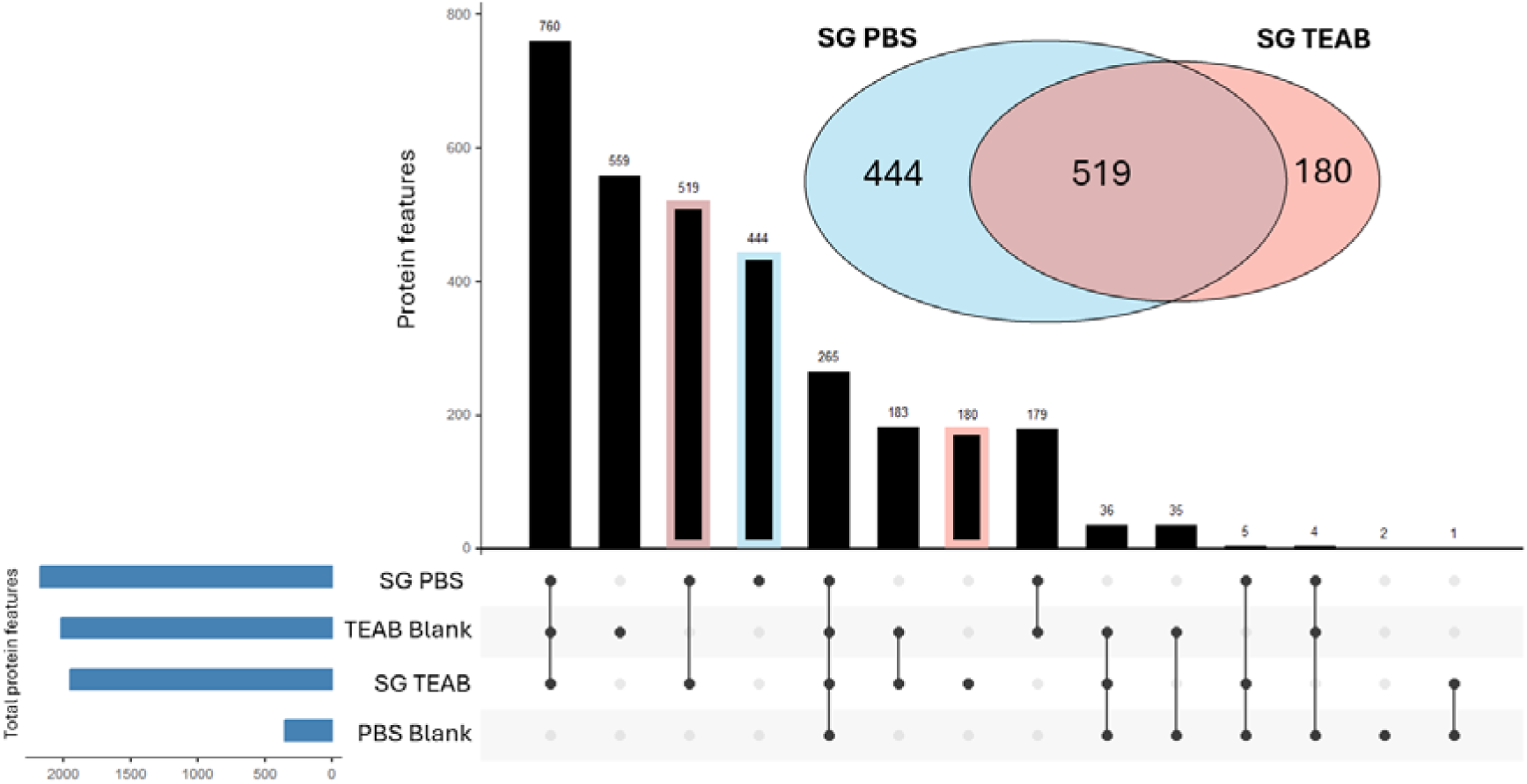
UpSet plot and Venn diagram showing overlap between proteins detected in SGs eluted in PBS or TEAB and corresponding blank controls. The highlighted column corresponds to proteins detected in at least one SG sample under both elution conditions and absent from both blanks that were retained, yielding a final set of 519 SG-associated proteins.

To assess whether the SG associated proteins identified here reflected any cytosolic contamination, the validated 519-protein dataset was compared against an independently derived cytosolic proteome using Data-Independent Acquisition localisation of proteins (DIA-LOP) of U2OS cells [45]. In this study, differential centrifugation was used to fractionate cells in to 11 subcellular fractions, each enriched in different subcellular components. After digestion to peptides, the proteins in each fraction were quantified using DIA-based label free quantitation. The computational workflow described in [53] was applied to classify proteins to different subcellular compartments. Classification using a support vector machine (SVM), yielded a ranked list of 1327 high confidence cytosolic proteins (see **SI Table 3**). Only 27 of the 519 SG associated proteins overlapped with this cytosolic dataset, indicating minimal contribution from proteins classified as cytosolic by abundance-based fractionation. This low overlap supports the specificity of the capillary-based sampling approach for enriching SG associated proteins rather than highly abundant cytosolic components.

Further analysis of detection frequency of these 519 high-confident SG-associated proteins revealed that a large proportion were detected in a limited number of individual SG samples, with progressively fewer proteins observed across multiple samples (see **SI Table 2** and **SI Figure 1**). This distribution is consistent with stochastic detection effects inherent to low-input proteomics [54], coupled with inter-SG heterogeneity [55]. When considered by elution condition, a greater number of proteins were detected exclusively in PBS-eluted SGs (444) compared to TEAB-eluted samples (180). Furthermore, PBS-eluted samples also exhibited a higher proportion of proteins detected across multiple SG samples, likely due to losses caused by bubbling of the TEAB samples during elution. These differences demonstrate how elution conditions can influence protein detection across SG samples.

To characterise functional relationships within the high-confidence SG proteome, the 519 SG associated proteins were analysed using the STRING database [50], which integrates known and predicted protein-protein associations and functional annotations. Functional enrichment analysis revealed significant representation of proteins involved in mitochondrial gene expression and mitochondrial translation, followed by pathways associated with rRNA processing, ribosome biogenesis, ribonucleoprotein complex biogenesis, and RNA metabolic processes (see **Figure 5A**). While the enrichment of RNA-associated pathways is consistent with established SG biology, the prominence of mitochondrial-associated processes raised the possibility of organelle carryover during subcellular sampling, although functional and physical cross talk between SG and mitochondria has been described before [48].

**Figure 5.**
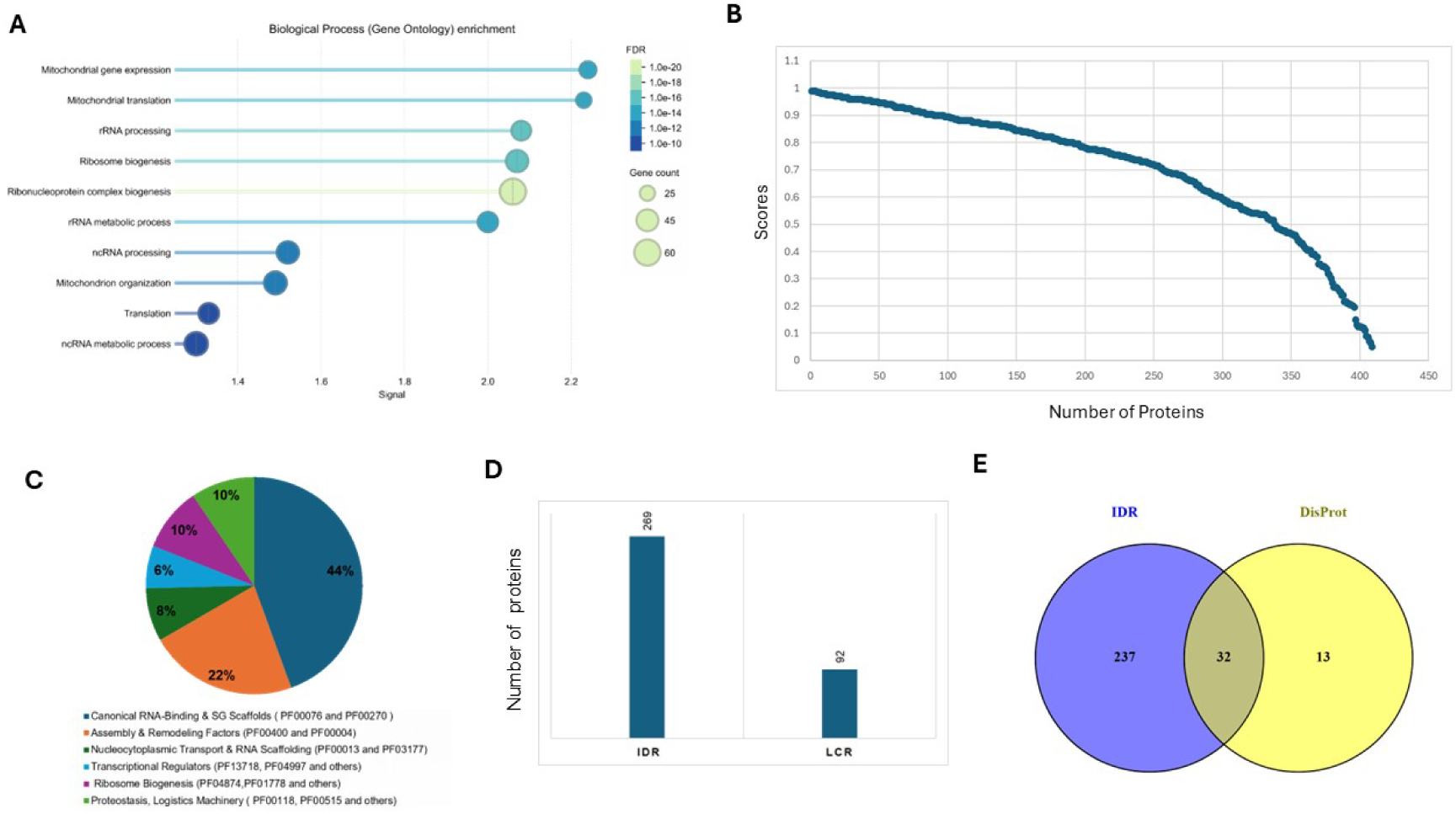
**A** STRING functional association network and pathway enrichment analysis of the 519 high-confidence SG associated proteins, with pathways ranked by STRING signal score. **B** Distribution of catGRANULE scores across the SG-associated proteome **C** Domain enrichment analysis of the refined SG proteome. **D** Bar chart illustrating the number of proteins containing predicted intrinsically disordered regions (IDRs) compared to those harbouring specific low-complexity regions (LCRs). **E** Venn diagram showing the overlap between computationally predicted IDRs and experimentally validated disordered regions curated in the DisProt database.

To address this potential carry over and focus subsequent analyses on the most SG-relevant candidates, organelle-associated proteins were excluded, yielding a refined set of 405 (see **SI Table 4**) proteins for downstream characterisation. Of these, 46 have been previously published as SG-associated proteins [56] [57] [58] (see **SI Figure 2** for full list and frequency distributions). These include RBPs such as FXR1, translation factors such as eIF3 subunits, and RNA helicases such as DHX30.

To assess whether the refined SG proteome retained the structural and functional characteristics expected of SG-associated proteins, we next examined the domain architecture of a representative subset of proteins (n = 63) spanning both previously reported and newly identified candidates. This subset was selected based on the presence of Pfam-annotated domains associated with RNA binding and condensate dynamics, enabling assessment of whether novel candidates share features of established SG components. Domain enrichment analysis revealed that 44% of proteins contained canonical RNA-binding and scaffolding domains, including RNA recognition motifs (RRMs) and RGG-rich regions, consistent with established SG components (see **Figure 5C**). A further 22% of proteins were associated with assembly and remodelling functions, including WD40-repeat proteins and AAA ATPases, which are implicated in regulating condensate dynamics and liquid–liquid phase separation (LLPS). The remaining 34% comprised proteins linked to proteostasis, ribosome biogenesis, and intracellular transport. Together, these findings indicate that both known and newly identified proteins integrate within the expected functional architecture of SGs, supporting the biological coherence of the refined dataset.

To characterize the biophysical properties of the identified SG proteome, we also employed the catGRANULE 2.0 algorithm to predict the liquid-liquid phase separation (LLPS) propensity of the 405 candidate proteins [59] (see **Figure 5B**). Our analysis revealed that approximately 60% of the identified proteins exhibit high catGRANULE scores, indicating a significant enrichment of features associated with condensate formation, such as structural disorder and specific amino acid composition and patterning (e.g., arginine/glycine-rich motifs). The observed distribution profile characterized by a sustained high propensity across most of the sequence is consistent with the ‘scaffold’ model of SG assembly, where a dense network of multivalent interactions drives the transition from a diffuse cytoplasmic state to a condensed liquid phase. Together, these findings provide strong computational support for the specificity of the isolated SG proteome and suggest that it is enriched in proteins with intrinsic capacity to participate in phase separation.

Using the MobiDB-lite consensus via UniProt, we also identified 269 proteins containing at least one intrinsically disordered region (IDR). We further screened these proteins for compositional bias and found that 92 proteins harboured significant sequence biases of low-complexity regions (LCRs) and clusters of acidic or basic residues (see **Figure 5D**). These structural signatures further confirm that these proteins possess the hallmark biophysical properties of SG components, consistent with previous studies [60]. Cross-referencing our candidates against the DisProt database identified 45 proteins with experimentally validated disordered regions. Matching these 45 proteins against our 269 predicted IDRs revealed that 32 proteins were captured by both datasets (see **Figure 5E**).

Notably, while 46 proteins within the refined dataset have previously been reported as SG-associated [58], the majority of identified proteins have not been described in SG proteome databases. This included protein families commonly associated with SGs such as DEAD-box helicases (e.g. DDX18 or DDX39B), stress-associated proteins (e.g. AHNAK2, BAG2) and metabolic enzymes (e.g. NUDT1). These findings indicate that the capillary-based sampling workflow combined with LC-MS proteomic analysis can identify additional candidate SG-associated proteins beyond those captured by bulk purification approaches.

To confirm these findings, the cellular distribution of some of these novel targets was validated using immunofluorescence following treatment with arsenite. As expected, the known SG residents present in our dataset FXR1 and DHX30 colocalised with both G3BP1 and eIF3E (known SG markers) in arsenite-stimulated U2OS cells (see **Figure 6**). Next, we tested whether proteins not previously associated with SG localisation and identified through our subcellular sampling analysis also colocalised with SGs. Upon arsenite stimulation AHNAK2, DDX39B, NUDT1 and FKBP2 all colocalised with G3BP1 and eIF3E in U2OS cells, confirming their status as SG resident proteins and the validity of our approach to deepen our knowledge of SG composition.

**Figure 6.**
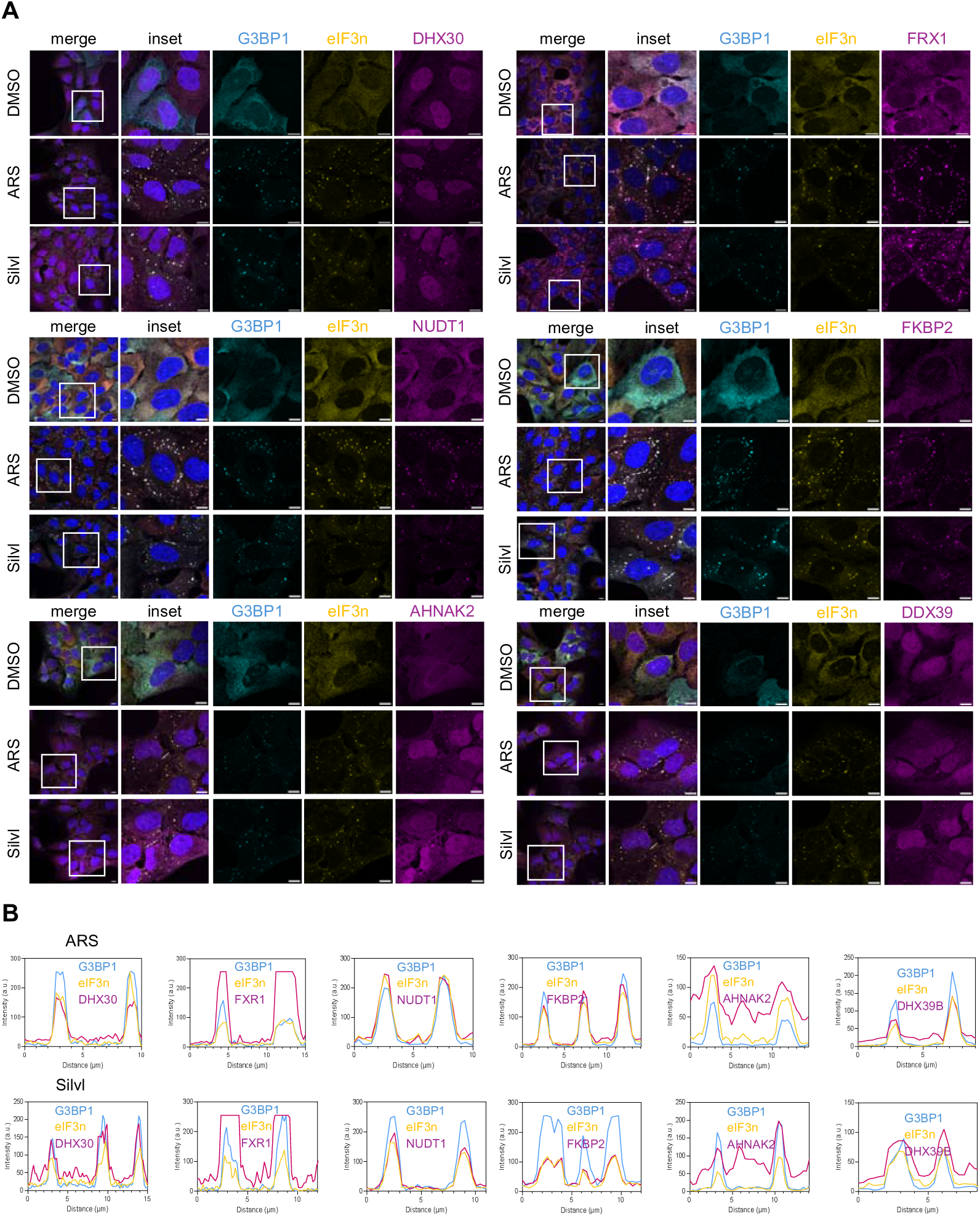
(**A**) U2OS were treated with 0.5 mM sodium arsenite (Ars) for 30 min, or 2 _μ_M silvestrol (Silvl) for 1 h, before harvest and cells were analysed by immunofluorescence for the canonical SG markers G3BP1 (cyan) and eIF3_η_ (gold), and resident proteins identified by mass spectrometry DHX30/FXR1/NUDT1/FKBP2/AHNAK2/DDX39B (magenta). Nuclei were stained with DAPI. Scale bars indicate 10 _μ_m. **(B)** Colocalisation of proteins was confirmed using line scan analysis in Image J for SGs marked by white lines in (A) inset panels.

Because SG assembled in responses to various stressors have been proposed to be compositionally different, we next asked whether these markers could be universal SG markers or specific to those assembled in response to arsenite treatment. SG assembly can be triggered following eIF2α-independent inhibition of translation, for example by blocking eIF4A or mTOR activity [61]. Thus, we examined AHNAK2, DDX39B, NUDT1 and FKBP2 cellular localisation following treatment with the eIF4A inhibitor silvestrol, a trigger for assembly of eIF2α-independent SGs [62]. Interestingly, all three markers redistributed to SGs upon silvestrol treatment, showing strong colocalization with eF3E and G3BP1. These results confirm both that our approach has the power to identify novel SG resident proteins and that AHNAK2, DDX39B, NUDT1 and FKBP2 are all potential core components of SGs. The conservation of SG localisation during both arsenite or silvestrol treatment also suggest these proteins act as core SG components rather than constituents of their dynamic shell with proposed stress-specific composition.

## Discussion

This study demonstrates a new method of capillary-based sampling coupled to LC-MS proteomics, which has enabled subcellular characterisation of a dynamic, membraneless organelle. Using this workflow, we recovered both previously reported and candidate SG-associated proteins, several of which were further supported by orthogonal validation using immunofluorescence microscopy. Importantly, the use of two parallel elution conditions with corresponding blank controls provided an internal framework for confidence filtering.

Microscopic heterogeneity is a defining feature of SGs, with a range of SG sizes and shapes observed in response to similar stressors within individual cells and across cell populations [63] [64]. Whether this results in functional differences or distinct stages of maturation during SG biogenesis remains an open question that subcellular sampling can start to address, for example through comparative analysis of SGs with differing physical properties.

Understanding SG composition is critical for dissecting their biological function. Here, we identify novel proteins associated with arsenite-induced SGs and show that these candidates accumulated in SGs formed in response to different stress pathways, including eIF4A inhibition-mediated translational shut-off. These findings suggest that the novel identified proteins may represent core SG components associated with the density rich phase of SGs, rather than dynamically shuttling in/out from SGs. Further work will be required to determine their roles during SG biogenesis and potential roles in SG maintenance or functions. Interestingly our study identified and validated AHNAK2 among the novel SG components. Previous work has demonstrated that AHNAK mRNA is regulated during cellular response and sequestered into SG upon acute ER or oxidative stress [65] and thus has been used as an mRNA marker for SGs. Our findings could suggest that both an mRNA and the protein it encodes are targeted to SGs, revealing co-regulatory process to both silence the expression of a transcript, and hijack its protein from the cytoplasmic milieu. Alternatively, given a recent work has proposed that SG can be hubs of localised translation, this could reflect local translation of the sequestered mRNA [66].

Compared with bulk SG purification methods, the capillary-based approach presented here enables sampling of live cells while retaining spatial information. This targeted sampling strategy preserves positional context and allows for the study of individual SGs rather than pooled populations, thereby capturing aspects of cellular heterogeneity that are inaccessible to bulk methods. Beyond SGs, this analytical framework is broadly applicable to other membranelles organelles, including P-bodies (PBs), nucleoli, and paraspeckles (PSPs). It also highlights the potential to perform proteomics analysis of other subcellular compartments (e.g. lysosome, mitochondria…) extracted from single cells by capillary sampling.

## Conclusion

In this study, we demonstrate the successful coupling of LC-MS proteomics with nanocapillary sampling for single-cell and subcellular analysis for the first time. This approach enables targeted extraction from live cells while preserving spatial context and avoiding bulk lysis or chemical fixation. By applying parallel elution conditions alongside corresponding blank controls, a high-confidence set of 405 SG-associated proteins were obtained, including 46 established in the literature and also previously unreported candidates. Localisation of selected novel candidates (AHNAK2, DDX39B, NUDT1 and FKBP2) was supported by immunofluorescence microscopy, and comparison with an independently derived cytosolic proteome further reinformed the specificity of the sampling approach. Collectively, these findings establish capillary-based subcellular sampling as a promising analytical strategy for defining organelle-resolved proteomes from single cells and provides new insight into the molecular composition of SGs.

## Acknowledgements

This work was supported by grants from Engineering and Physical Sciences Research Council (EPSRC) (EP/R031118/1, EP/X015491/1) and the Biotechnology and Biological Sciences Research Council (BBSRC) (BB/W019116/1; BBS/E/PI/23NB0003; BBS/E/PI/230002). The authors gratefully acknowledge Ed Emmott (University of Liverpool, UK) for his role in conceptualising this study. We also thank Harpreet Atwal (University of Surrey, UK), Jake Penny (King’s College London, UK) and Celia Jakob (The Pirbright Institute, UK) for their assistance with sample preparation.

## Data availability

The data that support the findings of this study are available from the corresponding author upon reasonable request.

