## Supplementary information for "Capillary-based Subcellular Sampling Uncovers the Stress Granule Proteome in Single Cells"

**Figures - Supplementary Information**

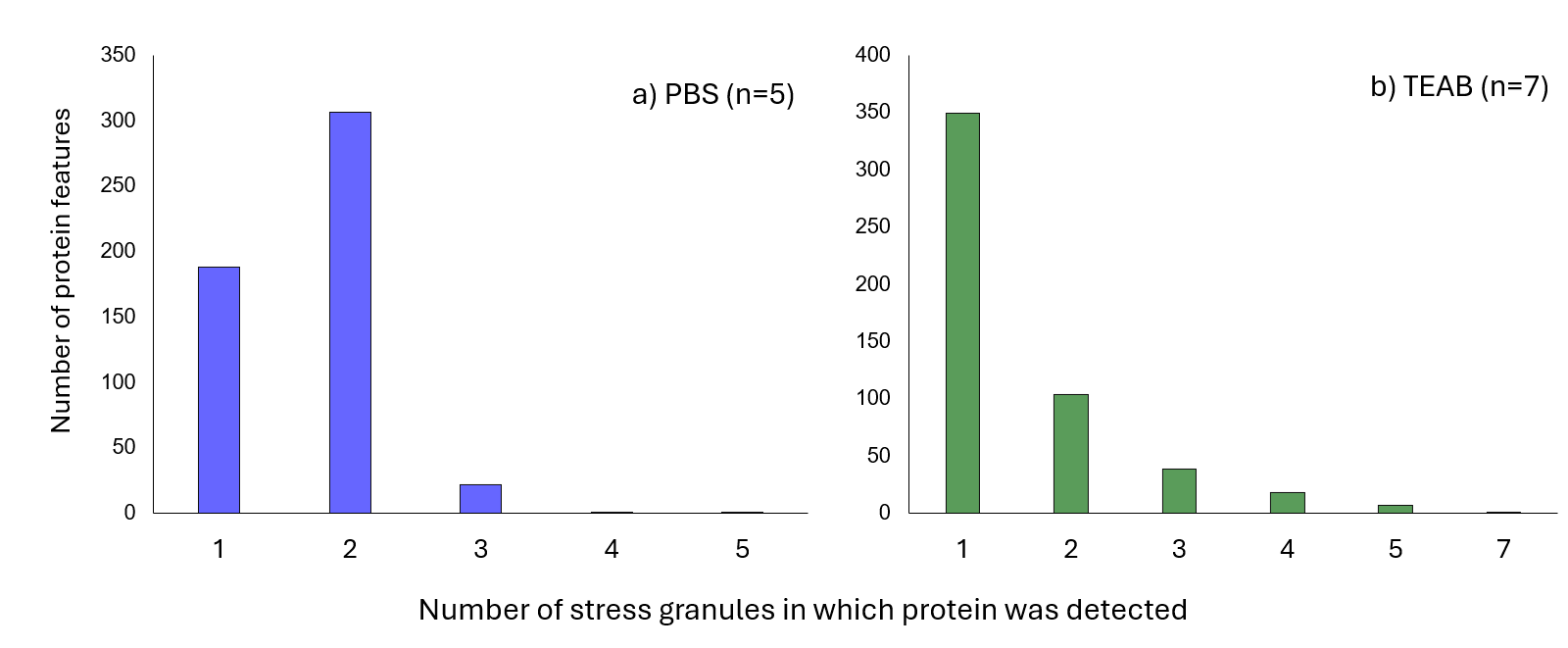

**Figure SI 1** Detection frequency of stress granule-associated proteins across independent stress granules eluted using **a)** PBS and **b)** TEAB.

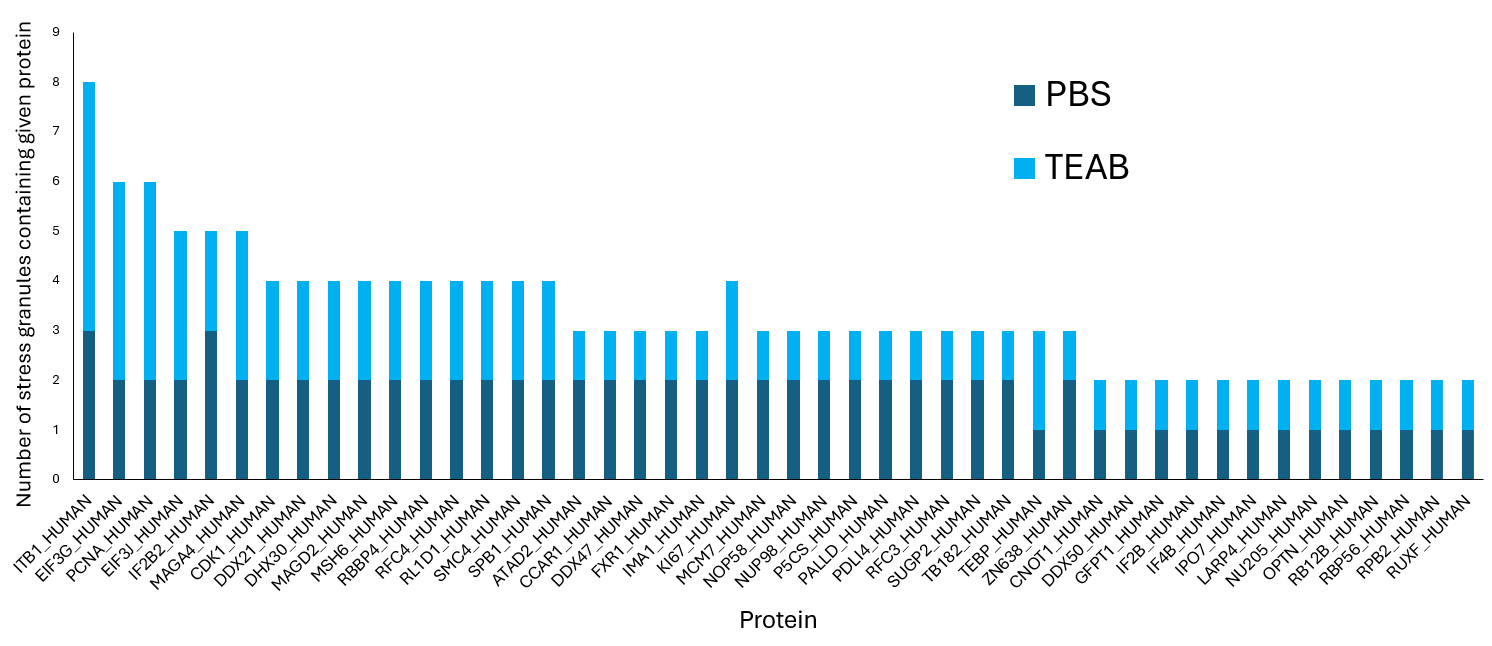

**Figure SI 2** List of 416 previously reported stress granule-associated proteins and detection frequency of the across independent stress granule samples eluted using PBS and TEAB.

**Tables - Supplementary Information (attached as .xlsx files)**

**Table SI 1** Table containing all proteins included in the volcano plot analysis, including log₂ fold change (log₂FC), raw p-values, −log10 transformed p-values, Benjamini–Hochberg false discovery rate (FDR)-adjusted p-values (padj), and significance classification. Proteins were considered significantly differentially abundant at *p* < 0.05 and |log₂FC| > 1. Upregulated proteins (green) have log₂FC > 1 (higher in Whole Stressed Single-Cells), and downregulated proteins (blue) have log₂FC < −1 (higher in untreated single-cell controls); proteins not meeting these thresholds were considered not significant.

**Table SI 2** Table containing the 519 high-confidence stress granule (SG)-associated proteins identified following filtering of the full dataset, and the SG samples in which each protein was detected. Proteins were retained only if detected in at least one PBS-eluted SG sample and one TEAB-eluted SG sample, while being absent from all PBS (n = 6) and TEAB (n = 5) blank controls.

**Table SI 3** Comparison of the high-confidence stress granule proteome with an independently derived cytosolic proteome. **(A)** Ranked list of 1327 high-confidence cytosolic proteins identified by DIA-LOP analysis of U2OS cells. **(B)** List of the 27 proteins overlapping between the 519 high-confidence SG-associated proteins identified in this study and the independently derived cytosolic proteome.

**Table SI 4** Refined high-confidence stress granule-associated protein dataset following exclusion of organelle-associated proteins.
